# Matrix Stiffness Dictates Doxorubicin-Induced Apoptosis by Modulating Cell-Cycle State in HeLa Cells

**DOI:** 10.1101/2025.11.11.687886

**Authors:** Natalie Calahan, Scott Burlingham, Ashok Prasad, Soham Ghosh

## Abstract

Drug resistance remains a major challenge in cancer treatment by contributing to recurrence and metastasis. Fractional killing, in which only a subset of cells undergo apoptosis after drug exposure, is a key contributor to this resistance and is influenced by genetic and nongenetic heterogeneity within the tumor microenvironment. Solid tumors display substantial variation in extracellular matrix stiffness, providing evidence that the mechanical context of cancer and stromal cells may play an important role in therapeutic response. Here, we investigated how substrate stiffness affects the dynamics of apoptosis and the mechanisms behind differences in the cell death response to doxorubicin (DOX). HeLa cells cultured on stiffer substrates exhibited enhanced caspase-3/7 activation and increased apoptotic cell death, whereas cells on soft substrates showed markedly reduced apoptotic signaling and improved survival. Although substrate stiffness altered cytoskeletal organization, pharmacological disruption of actin polymerization or actomyosin contractility did not influence nuclear DOX accumulation, indicating that cytoskeletal mechanics were not the primary factor in the stiffness-dependent sensitivity. Instead, flow cytometry revealed that substrate stiffness modulates cell-cycle distribution, with soft substrates enriched in the G1 population and a reduced fraction of cells in the DOX-sensitive S phase. Synchronizing cells at the G1/S phase boundary eliminated stiffness-dependent differences in apoptotic activation, demonstrating that cell-cycle state is a dominant driver of stiffness-mediated fractional killing. These findings highlight a mechanistic link between extracellular matrix mechanics and chemotherapeutic response by suggesting that microenvironment-regulated cell-cycle dynamics contribute to drug resistance in mechanically heterogeneous tumors.

## INTRODUCTION

In cancer treatment, one of the central challenges lies in the variability of cell responses to chemotherapeutic agents. When cells are treated with lethal drugs that induce apoptosis or other programmed death pathways, individual cells within a population show substantial heterogeneity in their response^1–4^. Even at maximal doses, many drugs fail to eradicate all cells due to the emergence of drug-tolerant subpopulations. These surviving cells can persist during extended drug exposure and regain proliferative capacity once the drug is removed, producing both drug-sensitive and drug-tolerant populations. This transient, reversible state is thought to contribute to the eventual development of stable drug resistance in vivo. Understanding the determinants of fractional killing is therefore important for identifying strategies that prevent the survival of these resistant cells.

Solid tumors often exhibit stiffness gradients, with softer cores and stiffer peripheral regions with biological implications in cells in different tumor regions^5–8^. This matrix heterogeneity introduces treatment challenges, as cell behavior and responses to therapy can be shaped by the mechanical properties of the microenvironment. Substrate stiffness has been shown to modulate drug response, and prior studies have reported differential chemotherapy sensitivity of cancer cells cultured on mechanically soft versus stiff substrates^9–11^. Cells also undergo characteristic changes in mechanical behavior and morphology in response to substrate stiffness. They maintain a level of internal pre-stress and modulate their stiffness to match that of their surroundings, a process known as stiffness-matching. On stiffer substrates and matrices, cells spread more and reorganize their actin cytoskeleton, whereas cells on soft substrates remain less spread and exhibit more fluid-like mechanical behavior^12,13^. Several fundamental processes, including proliferation rate and cell-cycle timing are also known to shift with substrate stiffness^14^. Because many chemotherapeutic agents preferentially target actively cycling cells, stiffness-dependent differences in cell-cycle progression may directly influence drug efficacy.

Despite increasing evidence that mechanical cues cause differences in therapeutic outcomes, the mechanisms through which substrate stiffness modulates chemotherapy response remains unclear. It is not known whether stiffness-dependent drug sensitivity arises from changes in cytoskeletal mechanics, differences in drug uptake and nuclear accumulation, or stiffness-driven alterations in cell-cycle distribution. Moreover, few studies have directly examined how mechanical cues influence early apoptotic commitment at the single-cell level, despite its central role in fractional killing.

The objective of this study is to determine whether substrate stiffness drives fractional killing of cancer cells and whether stiffness-dependent cell mechanics or cell-cycle state are behind this behavior. We exposed HeLa cells to doxorubicin (DOX) while culturing them on two-dimensional substrates of varying stiffness. We imaged and quantified activation of the apoptotic marker caspase-3/7 during DOX administration on a single-cell basis. Cytoskeletal-modifying drugs were used to assess whether stiffness-dependent differences in actin architecture influenced nuclear DOX accumulation. Flow cytometry and DNA content analysis were performed to determine whether substrate-driven changes in cell-cycle distribution contributed to variability in apoptotic sensitivity. Together, this approach allowed us to determine whether mechanics influence DOX response primarily through cytoskeletal pathways, drug transport, or stiffness-regulated shifts in cell-cycle progression.

## MATERIALS AND METHODS

### Cell culture

HeLa cells and HeLa cells stably expressing MY2A-GFP (HeLa MY2A-GFP) were cultured in High Glucose DMEM (Cytiva, SH30022.01) supplemented with 1mM sodium pyruvate (Gibco, 11360-070), 10% equafetal (Atlas Biologicals) and maintained at 37°C, 90% humidity and 5% CO_2_. TrypLE Express (Gibco, 12605-010) was used for passaging. For all live imaging experiments cells were maintained in complete culture medium. For all imaging experiments, 8 well μ-slides (ibidi, 80826) containing a polymer coverslip or thin PDMS coated coverslip were used, as described in the next section. For flow cytometry, cells were cultured in 100 mm petri dishes directly on tissue culture treated plastic or PDMS as described in the next section and reported in previous works^13^. All substrates were plasma treated and coated with bovine Type I collagen (Gibco, A1064401).

### PDMS gel formulation

Sylgard 527 (Dow Chemiclals, 01696742) and Sylgard 184 (Dow Chemicals, 04019862) were used according to manufacturer protocol. For this study, the soft gel refers to Sylgard 527, and the stiff gel is 8 parts Sylgard 527 thoroughly mixed with 1 part Sylgard 184 which results in the stiffness of 5 kPa and 80 kPa respectively, as reported in previous work. These formulations evenly spread on microscope cover glass (Fisher Scientific, 12541036), excess was allowed to run off allowing for a thin layer. Cover glass slides were place in a vacuum chamber for 20 minutes to remove all air bubbles. PDMS layer was then cured in an oven at 70°C for 2 hours. The glass slides were then attached to ibidi 8-well μ-slide frame with polymer bottom removed, using Sylgard 184 as a glue, and allowed to cure in the oven overnight.

### Drug treatment and cell cycle synchronization

HeLa My2A GFP cells were seeded at a density of approximately 20% a minimum of 24 hours before any drug treatment occurred. Cells were exposed to 5 ug/mL or 1.02 ug/mL of Doxorubicin (DOX) for varying time periods depending on the experiment. For disruption of cytoskeletal structure, pre-treatment of cells with Cytochalasin-D (Cyto-D) (0.4 μM) or Y27632 (Y27) (10 μM) lasted 24 hours before DOX application. Cell cycle synchronization was performed using a double thymidine block to arrest cells at the G1/S phase boundary by interrupting the deoxynucleotide metabolism pathway and halting DNA replication^15^. Thymidine (Millipore Sigma, T9250) was dissolved in DPBS to create a 2mM solution in complete cell culture media which was applied to cells for 14 hours, followed by a 9-hour wash and release period, before a second 14-hour application and release into DOX (5 ug/mL).

### Live imaging of apoptosis

For live cell imaging NucBlue Live Cell Stain ReadyProbes reagent (Invitrogen, R37605) was added to the ibidi plate wells (2 drops/ml) and maintained inside the incubator for 30 minutes at 37°C, 90% humidity and 5% CO2. After that the sample was placed in the confocal microscope stage (Zeiss LSM 980), DMEM containing NucBlue, NucView Caspase-3/7 Enzyme Substrates (2 μM), and DOX (5 ug/mL) replaced all existing media. NucView substrates (Biotium) consist of a fluorogenic DNA dye coupled to the caspase-3/7 DEVD recognition sequence. The substrate, which is initially non-fluorescent, penetrates the plasma membrane and enters the cytoplasm. In apoptotic cells, caspase-3/7 cleaves the substrate, releasing the high-affinity DNA dye, which migrates to the cell nucleus and stains DNA with fluorescence. Thus, NucView caspase-3/7 substrates are bifunctional, allowing detection of caspase-3/7 activity and visualization of morphological changes in the nucleus during apoptosis. Imaging was performed using the 10x objective with the 405 and 488 nm lasers. The sample was maintained at 37°C, 90% humidity and 5% CO_2_ during the imaging session using an incubation chamber. Imaging was performed every hour for 5 hours using an 8 by 8 tile setup with tile size 1414 by 1414 μm.

### Quantification of apoptosis and cell viability

Fluorescent intensity of caspase-3/7 activity for live cell imaging was quantified on a single cell basis using the ImageJ TrackMate plugin. The Difference of Gaussians (DoG) detection algorithm was used with an estimated average cell size of 30 μm. Cells were tracked using the simple Linear Assignment Problem (LAP) tracker with a linking max distance of 30 μm, gap closing max distance of 30 μm, and a gap closing max frame gap of 1. Tracking was based on the NucBlue fluorescence, with caspase activity intensity also recorded. Separately, sell cycle synchronized samples were fixed with 4% paraformaldehyde (PFA) following DOX application for accurate quantification of cell apoptotic activity at a specific timepoint and staining with NucBlue Live Cell Stain ReadyProbes reagent and NucView Caspase-3/7 Enzyme Substrates (2 μM). Caspase activity fluorescent intensity for fixed cells was quantified using the ImageJ threshold feature. Cell viability for all cases was performed using a live/dead assay kit (Thermo Fisher) consisting of Calcein AM and ethidium homodimer-1 according to the manufacturer’s protocol. Cells were imaged using a confocal microscope (Zeiss LSM 980) in the 488 nm (green) and 561 nm (red) channels using a 6 by 6 tile setup with tile size 1414 by 1414 μm.

### Quantification of F-actin fluorescence and DOX autofluorescence visualization

Cells were fixed with 4% PFA (in 1X PBS: Phosphate Buffer Solution) for 7 minutes at room temperature. Cells were permeabilized with 0.1% Triton X-100 in 1X PBS, washed with 1X PBS, and stained with GFP tagged phalloidin and DAPI. Imaging was completed using a 10x objective on a Zeiss LSM 980 microscope with 405 and 488 nm lasers. The fluorescent intensity of the F-actin was quantified using the threshold feature in ImageJ. To visualize DOX uptake into the nucleus, cells were exposed to DOX for 2 hours and fixed with 4% PFA. DOX exhibits autofluorescence when excited using a 488 nm laser. A confocal microscope (Zeiss LSM 980) with the 40x water objective was used to image cells with the 488 nm laser.

### Flow cytometry for cell cycle analysis

Cell cycle distribution was analyzed using an Attune NxT Acoustic Focusing Cytometer (Thermo Fisher Scientific). Cells were treated with DOX (1.02 ug/mL) for 12 hours prior to collection and resuspension in phosphate-buffered saline (PBS) containing Calcein AM (0.5 μM) (Thermo Fisher Scientific) to label live cells based on intracellular esterase activity and NucRed (1 drop/2mL) to label DNA. Cells were incubated for 30 minutes at 37 °C prior to flow cytometry. Samples were analyzed on the Attune NxT cytometer equipped with a 488 nm blue laser for Calcein AM (FITC channel; 530/30 nm filter) and a 561 nm yellow laser for NucRed (PE-Texas Red channel; 620/15 nm filter). Forward and side scatter parameters were used to exclude debris and aggregates, and doublets were removed by gating on pulse height versus area. A minimum of 10,000 live single-cell events were collected per sample. Data was analyzed using Attune NxT Software (Thermo Fisher Scientific) to demonstrate the proportion of cells in G□/G□, S, and G□/M phases among viable cells.

### Statistics

Parametric t-tests and p-value were used to quantify differences between the groups, as appropriate. Error bars in bar graphs represent the standard deviation. Number of technical replicates is reported in the figure captions.

## RESULTS

### Stiff substrates prime HeLa cells for increased Caspase-3/7 activation and apoptosis

Mechanical properties of the tumor microenvironment are recognized as regulators of cancer and stromal cell fate, but how matrix stiffness influences chemotherapeutic response remains poorly defined. To test whether substrate stiffness alters activation of apoptotic pathways and the fraction of cell death, we cultured cells on substrates spanning a range of stiffness, using plastic (∼1 GPa), a stiff PDMS (∼80 kPa), and a soft PDMS (∼5 kPa). Cells cultured on these substrates were then treated with doxorubicin (DOX), a topoisomerase II poison and DNA-damaging chemotherapeutic. Activation of apoptosis was monitored using a fluorescent reporter for caspase-3/7, the key executioner caspase. Representative images acquired every hour for 5 hours after DOX treatment revealed stiffness-dependent differences in caspase-3/7 activation (Fig. 1A). Cells on the soft PDMS exhibited minimal fluorescent signal, whereas cells on the stiff PDMS and plastic displayed robust caspase-3/7 activation, with the highest levels observed on the plastic substrate. Quantification of mean fluorescence intensity over time confirmed that caspase-3/7 activation remained low and nearly at the same level across time on soft PDMS, while stiff PDMS and plastic showed an increasing and significantly higher increase in apoptotic signal over the 5-hour period (Fig. 1B). To determine whether these population-level differences were driven by heterogeneous cell behavior, we performed single-cell tracking of caspase-/73 activation (Fig. 1C). On plastic, many cells showed rapid caspase-3/7 accumulation within 2–3 hours, but with considerable cell-to-cell variability. In contrast, soft and stiff PDMS produced much flatter single-cell traces; however, the stiff PDMS still demonstrates increased activation during the observation time. These findings suggest that substrate stiffness regulates the fraction of cells that commit to apoptosis instead of just shifting the mean level of caspase-3/7 expression.

**Figure 1:**
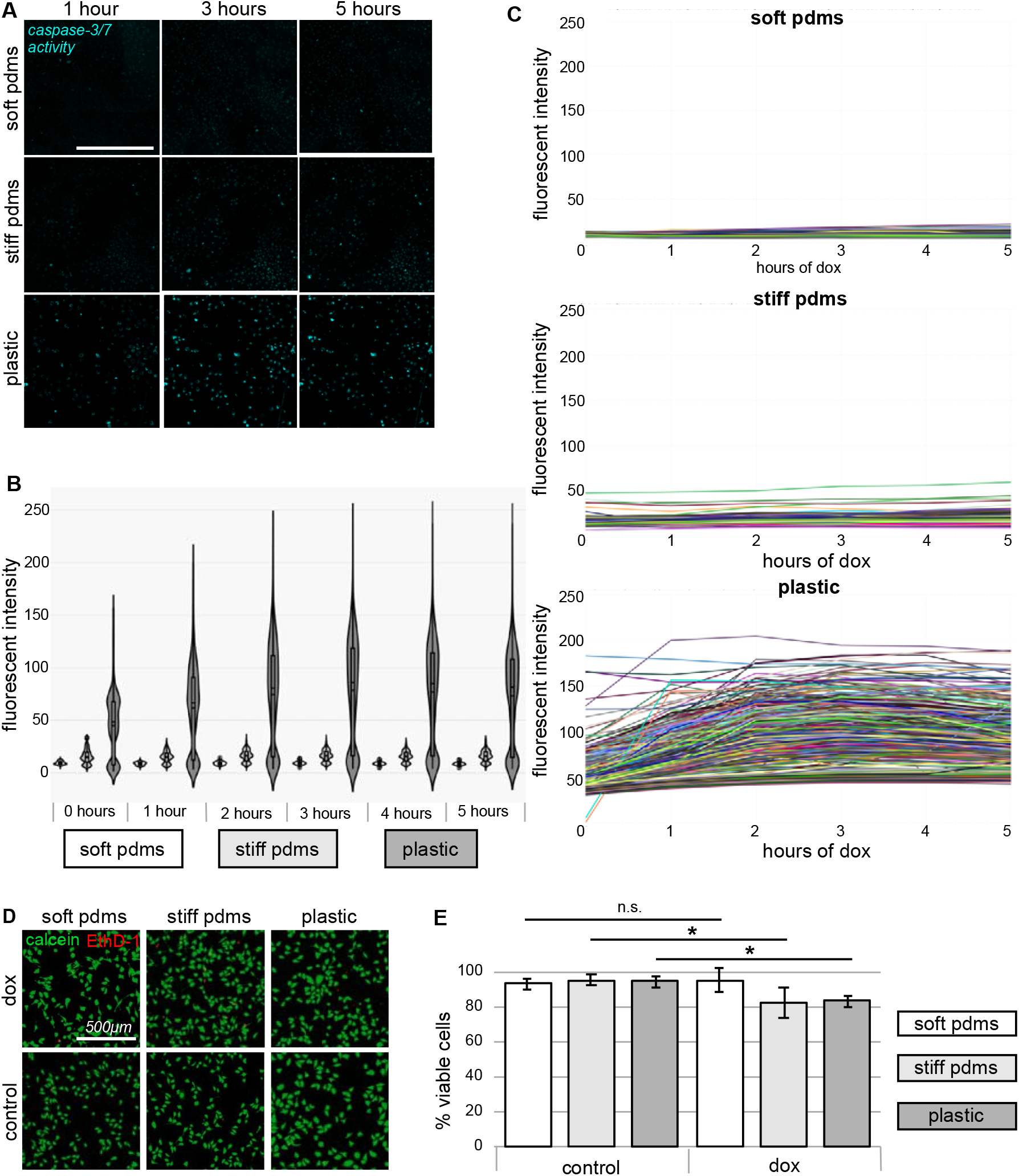
Impact of substrate stiffness on activation of apoptotic pathway and fractional killing in HeLa cells. **(A)** Represent ative fluorescence images of caspase-3/7 activity in cells cultured on substrates of increasing stiffness (soft PDMS, stiff PDMS, plastic) 5 hours after DOX treatment. **(B)** Violin plots of fluorescence intensity of caspase activity measured over a 5 hour time course shows stiffness-dependent increases in apoptosis activities. **(C)** Single-cell tracking of caspase-3/7 activity reveals heterogeneous apoptotic dynamics on stiffer substrates and minimal activation on soft substrates. **(D)** Live/dead staining 3 h after DOX treatment illustrates reduced cell death on soft PDMS compared with stiff PDMS or plastic. **(E)** Quantification of percentage of viable cells confirms significantly higher survival on soft hydrogels relative to stiffer conditions. *p<0.001. Data is based upon >1000 cells per group and 3 technical replicates.

We next tested whether the differences in the caspase-3/7 activity level translated into measurable effects of overall cell survival. Live/dead staining performed 3 hours after DOX exposure showed higher survival on soft PDMS relative to the stiff PDMS or plastic (Fig. 1D). Quantification confirmed stiffness-dependent protection from DOX, with over 95% viability on soft PDMS, and under 85% on stiff PDMS and plastic (Fig. 1E). Thus, a softer substrate not only reduced the apoptotic signaling but also decreased fractional killing. Together, these results demonstrate that substrate stiffness is a potent regulator of chemotherapeutic sensitivity with cells on soft matrices mostly evading caspase-3/7 activation and surviving DOX treatment, while cells on stiff substrates undergo higher rates of apoptosis and death.

### Cytoskeletal structural mechanics do not affect the nuclear accumulation of Doxorubicin

Based on our initial findings that substrate stiffness regulates apoptotic signaling and fractional killing (Fig. 1), we next sought to identify the upstream mechanisms that enable stiffness-dependent differences in chemotherapeutic response. For subsequent experiments, we focused on soft and stiff PDMS substrates rather than plastic because these conditions demonstrated a clear difference in cell survival following DOX treatment, and more closely reflect the physiologically relevant stiffness range of native and tumor□associated tissues. This approach allowed us to isolate relevant stiffness□dependent mechanisms without the additional effects introduced by ultra-rigid surfaces such as tissue culture plastic. This also allowed us to maintain similar autofluorescence of DOX through PDMS, which can be significantly changed when imaged through a polymer coverslip.

To test whether cytoskeletal organization was altered across these mechanical environments, we stained cells for F-actin using phalloidin. Quantification of F-actin fluorescent intensity revealed an increase in actin abundance on stiff PDMS, confirming that substrate stiffness drives cytoskeletal development (Fig. 2A). We next asked whether these stiffness-dependent cytoskeletal differences influence intracellular drug distribution, specifically nuclear uptake of DOX, which must accumulate in the nucleus act as a topoisomerase II poison. Autofluorescence imaging of cells revealed that DOX localized strongly to the nuclei of cells on stiff PDMS, whereas cells on soft PDMS displayed reduced and more heterogeneous nuclear signal intensity (Fig. 2B). To determine whether this difference was mediated by actin structure rather than the substrate stiffness alone, we pharmacologically disrupted the cytoskeleton on stiff PDMS using either Cytochalasin D (Cyto-D), which inhibits actin polymerization, or Y-27632 (Y27), a ROCK inhibitor that reduces actomyosin contractility. Neither of these treatments resulted in any distinct change to nuclear DOX accumulation, demonstrating that force generation through the actin structure was not required for DOX nuclear uptake.

**Figure 2:**
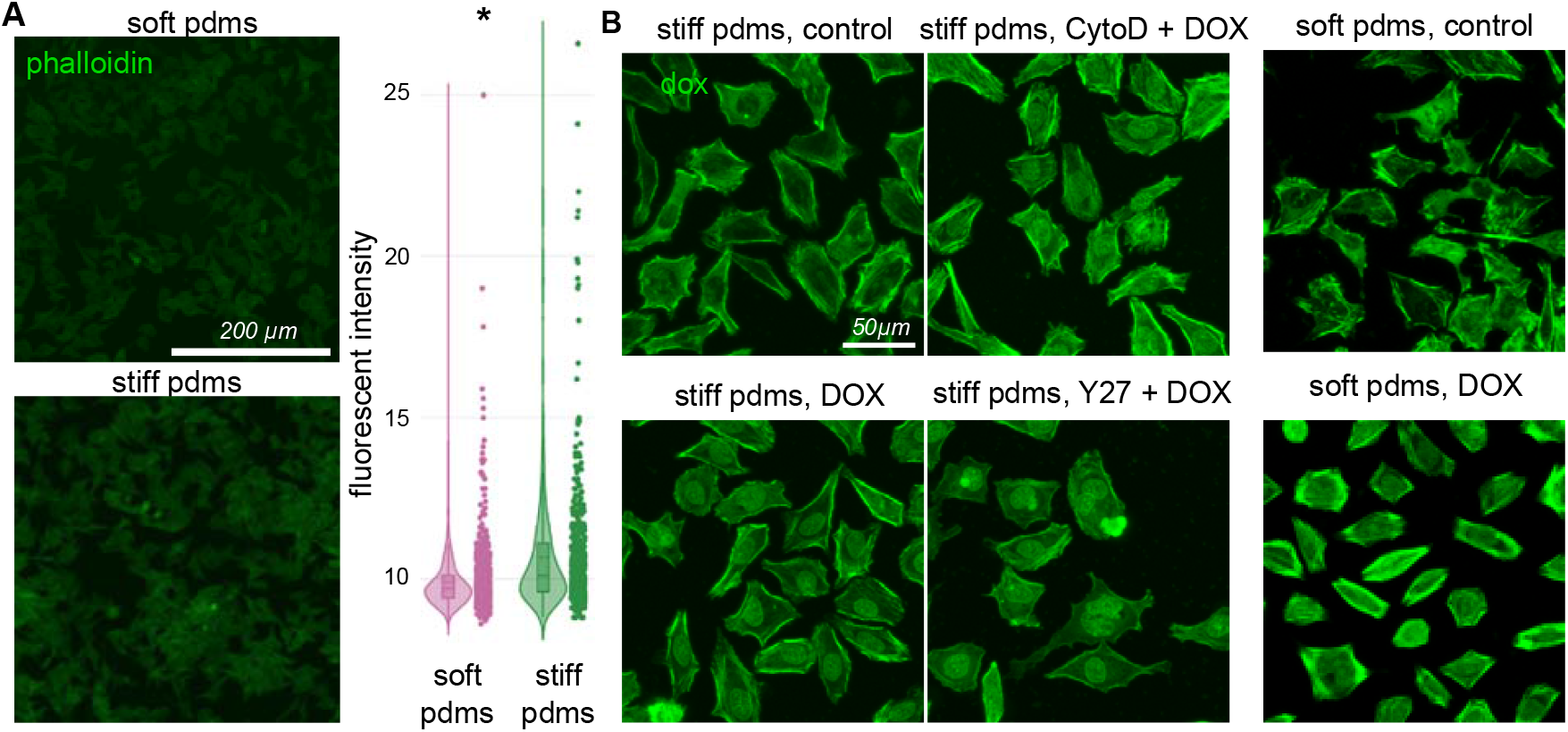
Role of cytoskeleton and substrate stiffness on DOX uptake and accumulation in HeLa cell nuclei. **(A)** Quantification of F-actin fluorescence intensity in cells cultured on soft versus stiff PDMS substrates demonstrates enhanced actin organization and cytoskeletal robustness on stiffer matrices. **(B)** Representative auto-fluorescence images of DOX-treated cells show nuclear drug accumulation under each condition. Cells on stiff PDMS exhibit significantly higher amount of nuclear DOX intensity compared to those on soft PDMS, and disruption of the actin cytoskeleton with Cytochalasin D (Cyto-D) or inhibition of ROCK-mediated cytoskeletal contractility with Y-27632 (Y27) do not reduce nuclear DOX accumulation on stiff substrates. *p<0.001. Data is based upon >500 cells/group.

These findings indicate that while increased substrate stiffness promoted formation of a robust actin cytoskeleton, it did not facilitate more efficient nuclear accumulation of DOX. In this case, cytoskeletal mechanics did not serve as a strong mechanistic link between matrix stiffness and chemotherapeutic response, leading to further studies into potential explanations for the stiffness-dependent apoptotic and viability outcomes observed in Fig. 1.

### Cell cycle progression due to substrate stiffness is a key factor that influences apoptosis dynamics

Given that disruption of the actin cytoskeleton did not eliminate stiffness-dependent differences in doxorubicin (DOX) nuclear accumulation (Fig. 2), we next asked whether the status of cell cycle state might also contribute to the differential fractional killing observed across substrate stiffness. DOX is a topoisomerase II–targeting drug whose efficacy is strongly linked to DNA replication. Therefore, if substrate stiffness alters the proportion of cells actively cycling through S phase, this could influence apoptotic sensitivity independently of drug uptake (Fig. 3A). We hypothesized that cells on soft substrates may survive because they are more likely to reside in drug□resistant phases of the cell cycle.

**Figure 3:**
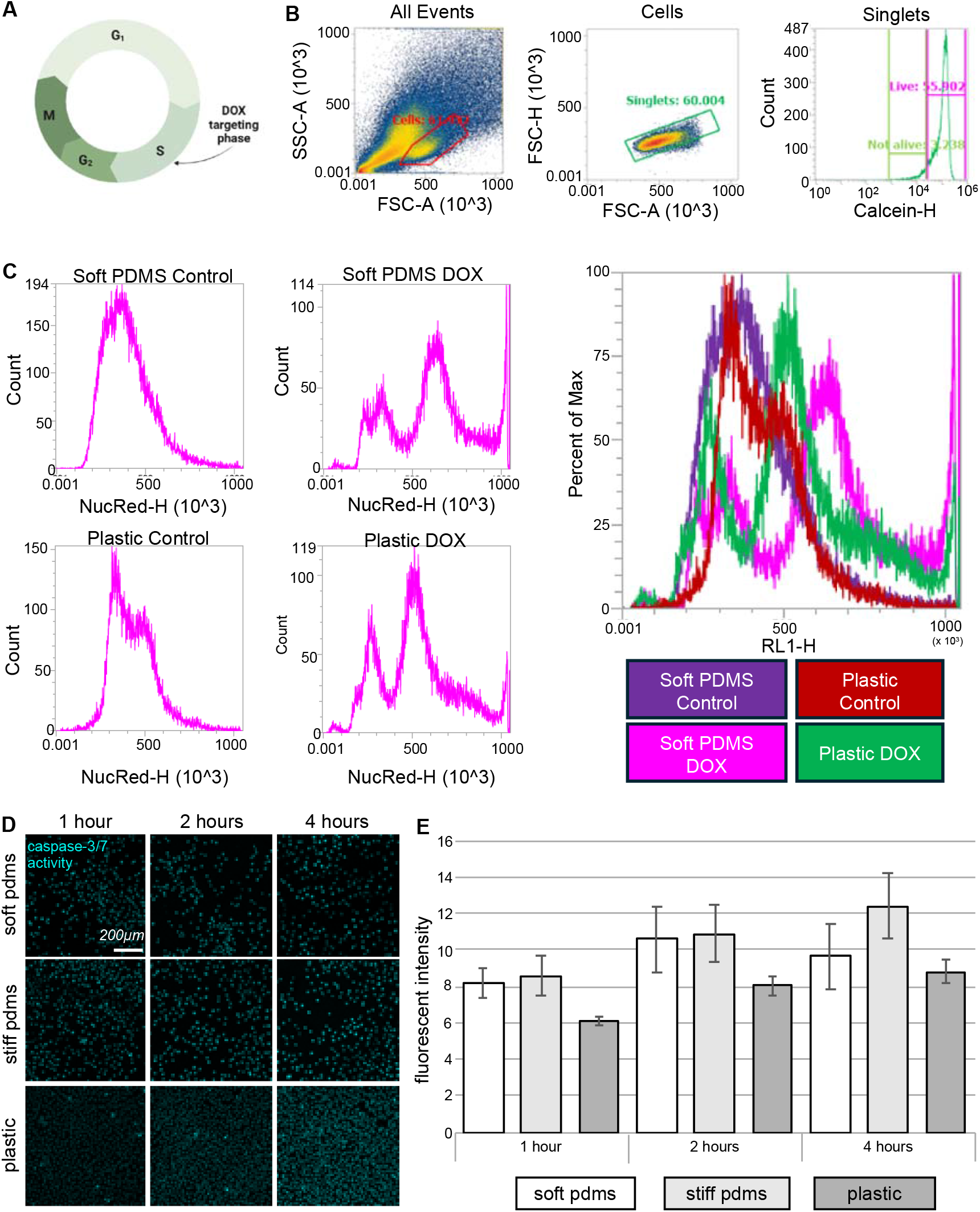
Effect of substrate stiffness dependent cell cycle progression on fractional killing. **(A)** Diagram of cell cycle phases showing where DOX targets the apoptotic pathway. **(B)** Plots of how cells were gated during flow cytometry to eliminate debris, doublets, and dead cells. **(C)** Flow cytometry analysis of DNA content showing the distribution of cells across G1, S, and G2/M phases on soft PDMS versus plastic substrates under asynchronous growth conditions. **(D)** Representative fluorescence images of caspase-3/7 activation in cells synchronized at the G1/S boundary prior to doxorubicin (DOX) treatment, demonstrating similar apoptotic activation on soft substrates compared with stiff substrates. **(E)** Quantification of mean caspase-3/7 activation over time following DOX treatment in synchronized cells shows that stiffness-dependent differences in apoptotic signaling are eliminated when cell cycle phase is controlled. Data is based on >10000 cells/ group.

To evaluate this, we first performed flow cytometry–based DNA content analysis (Fig. 3B) on cells grown under standard (asynchronous) conditions on soft PDMS versus plastic. Cells cultured on plastic displayed a larger fraction in S and G2/M, whereas cells on soft PDMS were enriched in G1 (Fig. 3C), indicating that substrate stiffness modulates proliferation dynamics and shifts the overall cell cycle distribution of the population. DOX-induced apoptosis is known to be most efficient in S phase, due to active DNA replication, and our results additionally confirmed this. After DOX treatment, cells on both substrates demonstrate a marked decrease in S-phase population, confirming this as the most susceptible phase.

To directly test this, we synchronized all cells to the G1/S temporal boundary prior to DOX treatment using a double thymidine block, thereby eliminating substrate-driven differences in baseline cell cycle composition. Representative caspase-3/7 activation images collected after DOX application showed that synchronized cells on stiff PDMS and plastic substrates still activated apoptosis, and synchronized cells on soft PDMS exhibited a similar caspase-3/7 response (Fig. 3D). Thus, synchronization equalized cell cycle phase at the time of treatment and eliminated substrate stiffness as an influence of apoptotic activation.

Quantification of caspase-3/7 fluorescence confirmed this pattern (Fig. 3E). Synchronization markedly reduced the difference between substrates compared to the asynchronous condition, with the plastic substrate showing even a lower activation compared to the PDMS substrates. These results indicate that distribution of cells in different cell cycle phases account for the stiffness dependent protection from DOX induced apoptosis seen on soft substrates.

## DISCUSSION

The mechanical properties of the tumor microenvironment are recognized as important non-genetic regulators of cancer cell behavior; however, their impact on chemotherapeutic response is not fully understood. In this study, we identified substrate stiffness as a key factor of doxorubicin (DOX) sensitivity, mainly acting through stiffness dependent shifts in cell-cycle distribution instead of cytoskeletal remodeling. Soft, and more compliant substrates reduced the caspase-3/7 activation and limited fractional killing by creating a protective state that allowed a larger fraction of cells to survive apoptotic induction through DOX exposure. Our experiments revealed a relationship between matrix stiffness and apoptotic pathway activation. Cells grown on stiffer substrates demonstrated increased caspase-3/7 activation after DOX treatment, while cell on the soft PDMS showed very little activation during this same time period. This difference in apoptosis progression also resulted in a difference in cell survival where most of the cells on the soft substrate remained viable compared to the decreased survival of those on stiffer substrates. These findings are consistent with previous studies which suggest that mechanical inputs can influence chemoresistance. However, most of that work focused on the activation of integrin-FAK survival pathways or YAP/TAZ signaling due to matrix stiffening^16,17^. The current study adds to this growing body of literature by showing that the protective effect of a soft matrix is linked with early apoptotic commitment over later survival related signaling.

We next asked how substrate stiffness led to changes in upstream regulators of apoptosis. Due to its known role in mechanotransduction we first looked at they cytoskeleton. As expected, cells on stiff substrates displayed an increased F-actin structure. We also observed higher nuclear accumulation of DOX under these conditions, raising the possibility that cytoskeletal tension could influence the drug transport. However, when actin polymerization or actomyosin contractility were inhibited, nuclear DOX levels remained relatively unchanged. This separates cytoskeletal mechanics from the DOX accumulation and suggests that while the actin network is responsive to stiffness, it is not the main mechanistic driver of the differing apoptotic response in these experiments. Although it is important to recognize that a cell’s nuclear pore complex is approximately 120 nm in diameter while the maximum diameter of a DOX molecule has been calculated to be around 1.5 nm, suggesting that DOX is capable of entering the nucleus regardless of differences in nuclear pore size. These findings align with reports showing that mechanical cues can regulate drug uptake and intracellular trafficking but also underscore that cytoskeletal tension is not universally required for these effects^18^.

Given the well-known dependence of DOX cytotoxicity on DNA replication, we next considered whether substrate stiffness influences the distribution of cell-cycle states within the population. Flow cytometry revealed that soft substrates increase the population of cells in G1, while stiffer matrices support larger S phase and G2/M fractions. Since DOX is most effective during active DNA replication, this shift provides a plausible explanation for why cells on soft PDMS survive treatment, as there are fewer cells in the drug vulnerable state. To directly test the possibility of such mechanism, we synchronized cells at the G1/S phase boundary. Once the populations of cells on all substrates were aligned in the same phase at the time of drug exposure, the stiffness-dependent difference in caspase-3/7 activation nearly disappeared. This provides evidence that cell cycle distribution is the primary factor governing stiffness dependent sensitivity to DOX. The mechanical environment does not only alter how cells handle stress but preconditions them by dictating whether they are even capable of engaging the apoptotic pathway at the time of drug administration.

While many studies report that stiff matrices promote drug resistance through pro-survival signaling of cancer stem cell programs, other work has shown that soft microenvironments can make cells less sensitive by placing them in slow-cycling or quiescent states^19,20^. Our results align with this second mechanism and help to detail previous contradictory observations across systems. Drugs that depend on DNA replication, including DOX, will naturally be less effective in environments that suppress proliferation. In contrast, other classes of chemotherapeutics may show differing patterns if their efficacy depends more on signaling pathways activated by stiff matrices. This context-dependent behavior highlights the need for careful consideration when drawing general conclusions about stiffness-induced resistance, especially as the magnitude of the effect may depend just as much on cell-cycle dynamics and drug class as on the mechanical cues themselves.

The implications of these can be inferred in the context of in vivo tumor environments. Many solid tumors contain regions of reduced stiffness due to ECM degradation, necrosis, or poor stromal organization. Cells within these niches may be more likely to be in slow-cycling states, making them less susceptible to chemotherapeutic agents that rely on active division. Our work suggests that the physical heterogeneity of the tumor microenvironment could contribute to spatially distinct drug responses and help explain why cancer often persists in softer and poorly structured regions of tumors.

This study has limitations in its examination of the variety of chemotherapeutic agents, as well as potential signaling pathways. Only one chemotherapeutic agent was used, and it is possible that other drugs with different mechanisms of action would engage mechanical signals through different pathways. Additionally, while cytoskeletal mechanics were ruled out as the main cause, upstream signaling that links substrate stiffness to cell cycle progression is yet to be determined. Mechanosensitive transcription factors such as YAP/TAZ, integrin-FAK signaling, and regulators of cyclins or CDK inhibitors are candidates for future studies.

Overall, we identified a mechanistic connection between extracellular matrix stiffness and chemotherapeutic response through which soft substrates protect cancer cells by biasing the population toward a protective cell cycle state. Our findings highlight the importance of mechanical context in therapeutic outcomes and suggest that targeting the mechanosensitive regulators of the cell cycle could create new opportunities to overcome microenvironment-driven chemoresistance.

## Acknowledgements

We acknowledge valuable discussions with Renzo Spagnuolo and Soumik Ghosh. The authors acknowledge the funding from NSF CMMI 2236710 and NSF CMMI 2227605.

## REFERENCES

1. Bertaux, F., Stoma, S., Drasdo, D. & Batt, G. Modeling dynamics of cell-to-cell variability in TRAILinduced apoptosis explains fractional killing and predicts reversible resistance. Plos Computational Biology 10, (2014).

2. Flusberg, D., Roux, J., Spencer, S. & Sorger, P. Cells surviving fractional killing by TRAIL exhibit transient but sustainable resistance and inflammatory phenotypes. Molecular Biology of the Cell 24, (2013).

3. Forcina, G., Conlon, M., Wells, A., Cao, J. & Dixon, S. Systematic quantification of population cell death kinetics in mammalian cells. Cell Systems 4, (2017).

4. Inde, Z., Forcina, G., Denton, K. & Dixon, S. Kinetic heterogeneity of cancer cell fractional killing. Cell Reports 32, (2020).

5. Paszek, M. et al./person-group>. Tensional homeostasis and the malignant phenotype. Cancer Cell 8, (2005).

6. Provenzano, P., Inman, D., Eliceiri, K. & Keely, P. Matrix density-induced mechanoregulation of breast cell phenotype, signaling and gene expression through a FAK-ERK linkage. Oncogene 28, (2009).

7. Chaudhuri, O. et al./person-group>. Extracellular matrix stiffness and composition jointly regulate the induction of malignant phenotypes in mammary epithelium. Nature Materials 13, (2014).

8. Mieulet, V. et al./person-group>. Stiffness increases with myofibroblast content and collagen density in mesenchymal high grade serous ovarian cancer. Scientific Reports 11, (2021).

9. Fan, Y. et al./person-group>. Substrate stiffness modulates the growth, phenotype, and chemoresistance of ovarian cancer cells. Frontiers in Cell and Developmental Biology 9, (2021).

10. Deng, B. et al./person-group>. Biological role of matrix stiffness in tumor growth and treatment. Journal of Translational Medicine 20, (2022).

11. Pan, H. et al./person-group>. Matrix stiffness triggers chemoresistance through elevated autophagy in pancreatic ductal adenocarcinoma. Biomaterials Science 11, (2023).

12. Solon, J., Levental, I., Sengupta, K., Georges, P. & Janmey, P. Fibroblast adaptation and stiffness matching to soft elastic substrates. Biophysical Journal 93, (2007).

13. Kaonis, S., Forman, J., Aboellail, Z. & Ghosh, S. High-throughput multiparametric quantification of mechanics driven heterogeneity in mesenchymal stromal cell population. Advanced Biology (2023).

14. Klein, E. A. et al./person-group>. Cell-cycle control by physiological matrix elasticity and in vivo tissue stiffening. Current Biology 19, 1511–1518 (2009).

15. Chen, G. & Deng, X. Cell synchronization by double thymidine block. Bio-Protocol 8, (2018).

16. Wei, S. C. et al./person-group>. Matrix stiffness drives epithelial – mesenchymal transition and tumour metastasis through a TWIST1 – G3BP2 mechanotransduction pathway. Nature Cell Biology 17, (2015).

17. Calvo, F. et al./person-group>. Mechanotransduction and YAP-dependent matrix remodelling is required for the generation and maintenance of cancer-associated fibroblasts. Nature Cell Biology 15, (2013).

18. Aydin, H. B., Ozcelikkale, A. & Acar, A. Exploiting matrix stiffness to overcome drug resistance. ACS Biomaterials Science and Engineering 10, (2024).

19. Bersini, S. et al./person-group>. Biomaterials A microfluidic 3D in vitro model for specificity of breast cancer metastasis to bone. Biomaterials 35, (2014).

20. Liukap, J. et al./person-group>. Soft fibrin gels promote selection and growth of tumorigenic cells. Nature Materials 11, (2012).

